# Muscle Mitochondrial Capacity and Endurance in Adults with Type 1 Diabetes

**DOI:** 10.1101/862086

**Authors:** Riley A. Hewgley, Bethany T. Moore, T. Bradley Willingham, Nathan T. Jenkins, Kevin K. McCully

## Abstract

The impact of type 1 diabetes (T1D) on muscle endurance and oxidative capacity is currently unknown.

**Purpose:** Measure muscle endurance and oxidative capacity of adults with T1D compared to controls.

**Methods:** A cross-sectional study design with a control group was used. Subjects (19-37 years old) with T1D (n=17) and controls (n=17) were assessed with hemoglobin A1c (HbA1c) and casual glucose. Muscle endurance was measured with an accelerometer at stimulation frequencies of 2, 4, and 6 Hz for a total of nine minutes. Mitochondrial capacity was measured using near-infrared spectroscopy after exercise as the rate constant of the rate of recovery of oxygen consumption.

**Results:** T1D and control groups were similar in age, sex, height, and race. The T1D group had slightly higher BMI values and adipose tissue thickness over the forearm muscles. Casual glucose was 150±70 mg/dL for T1D and 98±16 mg/dL for controls (P=0.006). HbA1c of T1D subjects was 7.1±0.9% and 5.0±0.4% for controls (P<0.01). Endurance indexes at 2, 4, and 6 Hz were 94.5±5.2%, 81.8±8.4%, and 68.6±13.5% for T1D and 94.6±4.1%, 85.9±6.3%, and 68.7±15.4% for controls (p = 0.97, 0.12, 0.99, respectively). There were no differences between groups in mitochondrial capacity (T1D= 1.9±0.5 min^−1^ and control=1.8±0.4 min^−1^, P=0.29) or reperfusion rate (T1D= 8.8±2.8s and control=10.3±3.0s, P=0.88). There were no significant correlations between HbA1c and either muscle endurance, mitochondrial capacity or reperfusion rate.

**Conclusions:** Adults with T1D did not have reduced oxidative capacity, muscle endurance or muscle reperfusion rates compared to controls. HbA1c also did not correlate with muscle endurance, mitochondrial capacity or reperfusion rates. Future studies should extend these measurements to older people or people with poorly-controlled T1D.

## INTRODUCTION

Type 1 diabetes (T1D) is a prevalent autoimmune disease resulting from specific immune-mediated destruction of pancreatic beta cells, which is managed using multiple daily injections of insulin or an insulin pump along with careful monitoring of blood glucose levels (1). If unmanaged or poorly treated, this disease can have consequences on all organ systems, especially cardiovascular and renal, due to impacts on macro- and microvascular systems, including atherosclerosis, endothelial permeability, and thickening of capillary walls (2). T1D is also associated with decreased mitochondrial oxygen consumption and impaired oxidative phosphorylation efficiency (3). Declining heart health in those with T1D could be a direct result of chronic deterioration of mitochondrial function due to oxidative stress (4, 5). People with T1D also report elevated fatigue that may be due to increased pain, sleep disturbances, depressive symptoms, or the physiological impacts of their condition (6). In a previous study, mitochondrial DNA mutations were observed in diabetic tissues potentially due to oxidative stress (7). A loss of muscle mass and fiber atrophy has been associated with T1D (8) as well as decreased muscle strength (9), but previous studies have been unable to confirm a relationship between fatigue and glucose control in people with T1D. Further evidence is needed to determine the onset and effects of T1D on mitochondria function and implications for muscle endurance.

Neuromuscular electrical stimulation (NMES) is a technique that has been used in combination with a tri-axial wireless accelerometer to determine skeletal muscle endurance (10–12). NMES has also been used in various other forms of assessing fatigue including handgrip fatigue (13) due to reduced variability associated with controlled, artificial exercise. Near infrared spectroscopy (NIRS) has been used in previous studies as a non-invasive approach to measuring muscle oxygen consumption as a gauge of mitochondrial capacity (14–16) as well as skeletal muscle blood flow (17). These technologies have been applied to other populations, such as those with Friedrich’s Ataxia (10), spinal cord injuries (18), and peripheral vascular disease (19).

The purpose of this study was to determine the association of T1D in young adults with muscle endurance, mitochondrial capacity, and muscle reperfusion. We hypothesized that T1D would be associated with reduced muscle endurance, a decreased rate of recovery of muscle oxygen consumption, and a longer reperfusion rate.

## METHODS

### Participants

Subjects either had a diagnosis of T1D (n = 17), or were healthy controls (n = 17) and between 18-40 years old were recruited to produce similar group averages for age, sex, height, and weight. Exclusion criteria included the presence any neuromuscular, cardiovascular, and endocrine diseases (except for T1D). The study was approved by the University of Georgia IRB, and all subjects gave informed consent.

### Experimental design

This experiment was a cross-sectional evaluation between T1D and control groups. After eligibility was determined with a screening form, subjects completed a medical history. Testing sessions consisted of three main components: clinical measurements, an electrical stimulation endurance test, and a near-infrared spectroscopy (NIRS) test of mitochondrial capacity. The tests were performed on the wrist-finger flexors of the non-dominant forearm: the flexor carpai radialis, flexor carpai ulnaris, and palmaris longis muscles. The forearm muscles were chosen because they are a relatively untrained group of muscles that are not directly influenced by regular physical activity levels.

### Experimental Procedures

Health indicators included measurements of casual blood glucose, HbA1c, and adiposity above the forearm muscles. HbA1c and casual glucose measurements were taken for every participant. Casual glucose was taken by finger-stick at the time of testing without prior instruction or fasting, and measured with a glucometer (OneTouch LifeScan, Inc., Milpitas, CA). HbA1c was measured by finger-stick in the same manner with a rapid HbA1c device (Accu-chek, Roche, Indianapolis, IN). Adipose tissue thickness over the forearm muscle was measured with B-mode ultrasound (GE Logic Q, GE Medial, Milwaukee, WI) as a routine control variable for NIRS measurements. This measurement was recorded and used to adjust separation distances of the NIRS optical probes.

The endurance test (12) was performed as the participant lay supine on a padded table with the non-dominant arm outstretched comfortably and secured at 90 degrees on a table extension to decrease movement (Figure 1a). A tri-axial, wireless accelerometer (WAX9, Axivity, UK) was attached to the skin at the belly of the forearm muscles using double-sided adhesive tape. Two electrode pads (2in × 4in) were placed 2 cm proximal and 2 cm distal of the accelerometer. Prior to the start of the test, stimulation current (Theratouch 4.7, Rich-Mar, USA) was adjusted until a characteristic, visible contraction was observed. Data collection began with a 15-second baseline measurement, and then stimulation of the forearm muscle began by applying twitch electrical stimulation in three stages of increasing stimulation frequencies (2, 4, and 6 Hz). Each stage was 3 minutes, was separated by 15 seconds of rest, and concluded with 15 seconds of recovery after stimulation. Twitch-induced surface oscillations in the X, Y, and Z directions gave the force of muscle contractions measured by the accelerometer. Data were collected at an acquisition frequency of 400Hz via a Bluetooth communication port. Reproducibility for the 2,4, and 6 Hz endurance index measurements in control subjects was 2.5-7.4% (12).

**Figure 1.**
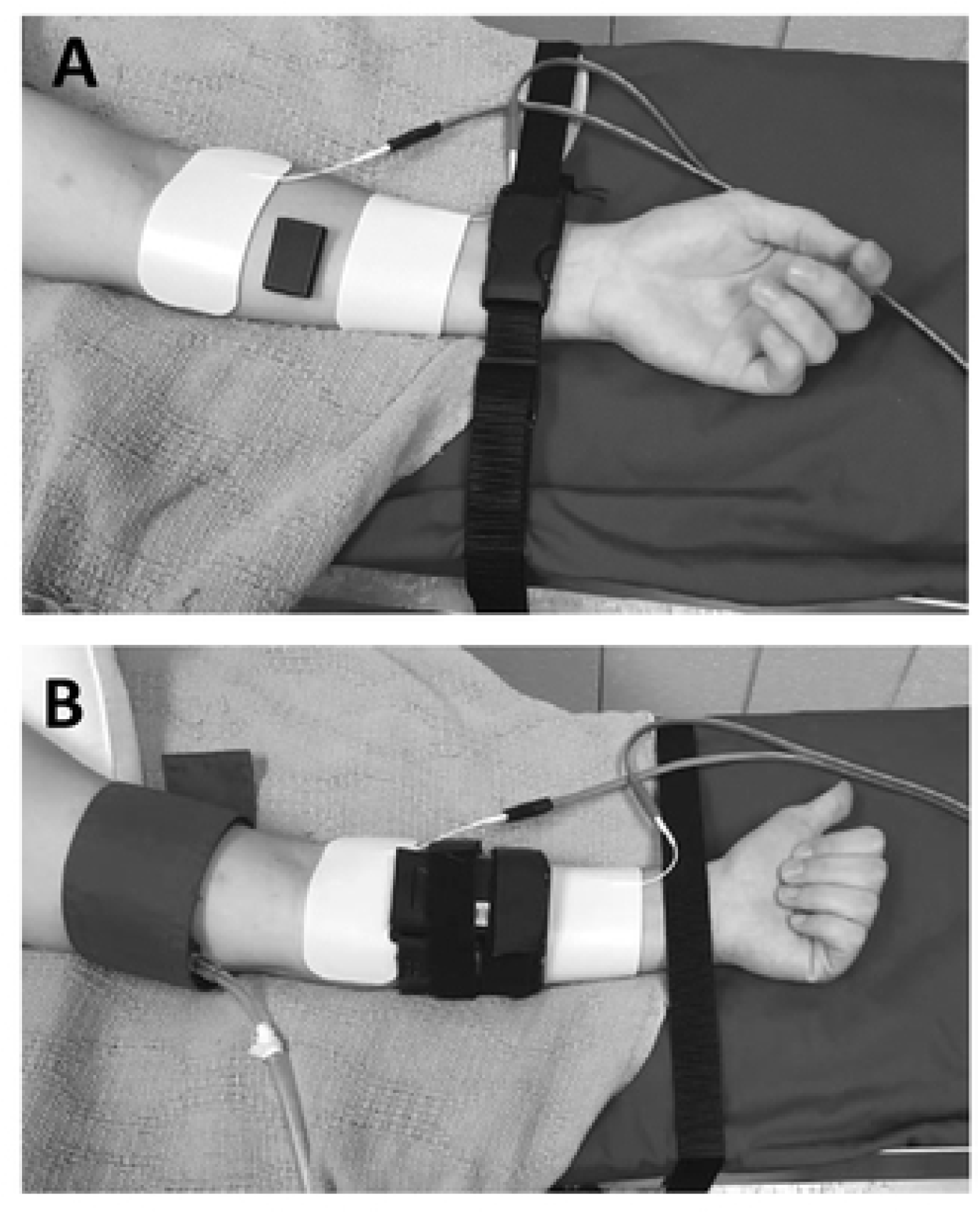
A) Experimental set up for the measurements of muscle endurance on the forearm muscles. B) Experimental set up for measurements of muscle mitochondrial capacity and muscle reperfusion rate. The white pads are stimulation electrodes. The black objects between the electrodes are either the accelerometer or the NIRS device. The blue wrapping is the cuff for the rapid cuff inflator.

**Figure 2.**
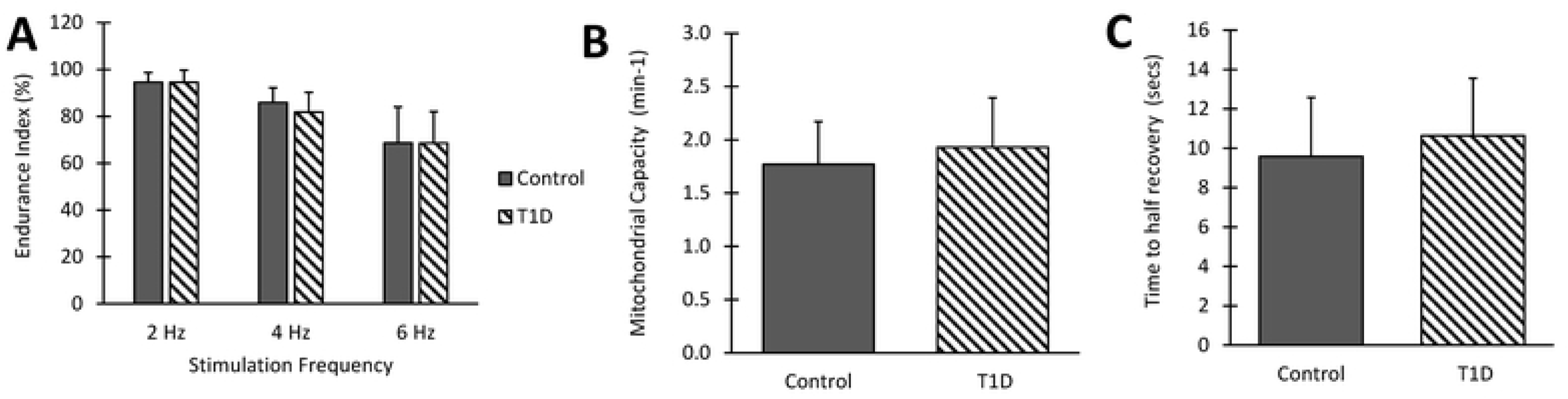
A) Muscle endurance measured in percentage in people with T1D versus controls at 2, 4, and 6 Hz. n=17 both groups. B) Mitochondrial capacity given in rate constant (1/min) in people with T1D versus controls. n=17 both groups. C) Muscle reperfusion rate measured in seconds in people with T1D versus controls. n=17 both groups. Values are means ± SD. No statistical differences were found for any of the comparisons in these figures.

Mitochondrial capacity (20) was measured as the rate of recovery of oxidative metabolism after exercise using NIRS (Figure 1b). The NIRS device (Oxymon, Artinis ltd, Amsterdam, NL) was placed directly above the belly of wrist-finger flexors. Electrical stimulation pads were placed on the forearm, above and below the NIRS device, and an inflatable cuff was placed above the elbow proximal to the NIRS device. The inflation pressure of the cuff was approximately 100 mm Hg above systolic pressure. After 30 seconds of rest, the cuff was inflated for five minutes or until plateau of the oxygenated and deoxygenated signals to deplete the oxygen in the muscle by occluding arterial blood flow, representing 0% O_2_ saturation or ischemia. Upon release of the cuff, oxygen returned to the muscle, creating a hyper-oxygenation response representing 100% O_2_ saturation or hyperemia. These two points were used in the calibration of the signal for the NIRS signals. The time from the release of the cuff to 100% O_2_ saturation was recorded as the time-to-half recovery, or reperfusion, of oxygen saturation and was used as an index of microvascular blood flow (21). After the calibration cuff, the muscle was stimulated for 15 seconds at 6 Hz as exercise to increase metabolic rate. Immediately following electrical stimulation, a recovery test cuff protocol was performed: cuffs 1 through 10 were 5 seconds on, 5 seconds off; cuffs 11 through 15 were 7 seconds on, 7 seconds off; cuffs 16 through 18 were 10 seconds on, 15 seconds off, and cuffs 19 and 20 were 10 seconds on, 20 seconds off. The exercise and cuff protocol was repeated, and the two recovery tests were averaged to measure the rate of recovery of muscle oxygen consumption, representing mitochondrial oxidative capacity. Reproducibility of the mitochondrial capacity measurements has been reported to be 12% in a previous study on the medial gastrocnemius muscle (22).

### Data Analysis and Statistics

For the muscle endurance test, peak-to-peak oscillations throughout stimulation was analyzed through Microsoft Excel and a MatlLab analysis routine (12). The endurance index was calculated as the end acceleration of the frequency period divided by the peak acceleration at the start of the stimulation period multiplied by 100 percent to allow for comparison within and between groups. Oxymon software was used to collect the data for the NIRS test, and the data was analyzed with a Matlab analysis routine. Comparisons between groups were made with unpaired, two-tailed T tests using SPSS. Statistical significance was accepted with a p value <0.05. Power estimates suggested that with a sample size of 17 per group and a population variance of 20%; a difference between groups of 20% could be detected in the mitochondrial rate constant with a power of 0.83.

## RESULTS

The subject’s physical characteristics are shown in Table 1. Subjects with T1D had higher casual glucose and HbA1c levels than controls, as expected. The subjects with T1D also had slightly higher BMI and forearm adiposity measurements.

**Table 1.** Characteristics of the Control and T1D groups.

|  |  | **Control** | **T1D** | **p value** |
| --- | --- | --- | --- | --- |
| **Age** | **Years** | 23.2 ± 4.1 | 23.7 ± 5.8 | 0.76 |
| **Sex** | **Male** | 3 | 6 |  |
|  | **Female** | 2 | 11 |  |
| **Race** | **White** | 16 | 16 |  |
|  | **Black** | 1 | 1 |  |
| **Height** | **cm** | 169.0 ± 9.6 | 170.4 ± 11.1 | 0.69 |
| **Weight** | **kg** | 67.8 ± 14.7 | 76.8 ± 13.0 | 0.07 |
| **BMI** | **kg/m<sup>2</sup>** | 23.6 ± 3.2 | 26.3 ± 2.7 | 0.01* |
| **Adipose** | **cm** | 0.4 ± 0.1 | 0.5 ± 0.2 |  |
| **Casual glucose** | **mg/dL** | 98 ± 16 | 150 ± 70 | 0.01* |
| **HbA1c** | **%** | 5.0 ± 0.4 | 7.1 ± 0.9 | < 0.001* |

n=17 both groups. Values are means ± SD. Asterisks (\*) indicate significance between all T1D and all controls.

Muscle endurance, measured as the endurance index, was not different between subjects with T1D and controls (p = 0.97, 0.12, 0.99 for 2, 4, and 6 Hz respectively) (Figure 1A). Muscle mitochondrial capacity measured as the recovery rate constant was also not different between subjects with T1D and controls (p = 0.29) (Figure 1B). Muscle reperfusion rate measured as the half time of recovery of oxygen saturation was not different between subjects with T1D and controls (p = 0.88) (Figure 1C). There was no relationship between HbA1c levels and Muscle endurance (6 Hz), mitochondrial capacity or muscle reperfusion rate (r^2^ values of 0.01, 0.10, and 0.01, respectively). There was also no relationship between subject age and time since diagnosis and the measurements of Muscle endurance (6 Hz), muscle mitochondrial capacity and muscle reperfusion rate (r^2^ < 0.1 for all comparisons).

## DISCUSSION

The primary findings of this study were that muscle endurance and muscle mitochondrial capacity were not different between adults with TID and controls. Previous studies have suggested that type 2 diabetes is associated with increased fatigue (reduced endurance) (23–27). However, there has been less research on muscle endurance and muscle mitochondrial capacity in people with T1D. Approximately 75% of people with T1D report persistent symptoms of fatigue (6), and T1D is thought to accelerate muscle aging (28) and result in pathological changes in skeletal muscle (8). One previous study by Cree-Green et al. reported reduced muscle mitochondrial OxPhos after exercise, marker of mitochondrial capacity, in adolescents (mean age of 15 yrs) with T1D (4). It is not clear how to compare our results to theirs. Cree-Green et al. reported no difference in phosphocreatine recovery rates after exercise, a marker of mitochondrial capacity analogous to the NIRS based recovery rate constants used in our study (29). In a study on adults with T1D during a two day endurance cycling event, investigators found significantly decreased power output was associated with low blood glucose levels, likely resulting from suboptimal dietary carbohydrate intake rather than a mitochondrial deficit (30). A study by Harmer et al. reported that any fatigue associated with T1D is not associated with impaired muscle calcium kinetics and that fatigue reported in people with T1D may originate outside the skeletal muscle (31). Our findings add to this small body of research, providing direct support for the hypothesis that T1D in adults is not associated with reduced muscle endurance or muscle mitochondrial capacity.

There were no differences muscle reperfusion rate between adults with T1D and controls. We had hypothesized that T1D might damage blood vessels due to chronic hyperglycemia and accompanying increased variability in blood glucose levels (32, 33). Previous studies have shown that the rate of muscle reperfusion is an indicator of impaired circulation (34, 35). While the subjects in our study were relatively young and had well-controlled blood sugar and HbA1c levels, previous studies have suggested vascular abnormalities in similar populations (36, 37). Additional studies on older subjects with T1D or in subjects with a history of poor glycemic control would are needed to better understand the impact of T1D on muscle reperfusion rates.

In the present study, no correlations were found between well-controlled people with T1D in the parameters tested regarding age, time since diagnosis, or HbA1c levels. Previous studies have also found weak or no relationships within populations of people with T1D and HbA1c levels or other indications of glycemic control (37). However, some studies have reported relationships between vascular function (nitrate stimulated brachial artery diameter changes) and HbA1c (33). The inconsistency of the relationship between vascular dysfunction and glycemic control in people with T1D may reflect the impact of compounding health risks, such as obesity, inactivity, stress and genetic factors (38, 39).

The strengths of this study were that the tests performed were non-invasive and time-efficient (40). Even vulnerable populations such as those with spinal cord injury are able to tolerate NIRS testing and electrical stimulation (41). These methods and technologies can easily be used in a clinical setting to evaluate the individual with little inconvenience and a high level of accuracy and reproducibility (20). There are alternative ways to measure mitochondrial capacity including ^31^P magnetic resonance spectroscopy (42); however, this approach is limited by its high cost and limited availability. In addition, there are complications related to strong magnetic fields used by P magnetic resonance spectroscopy and the use of insulin pumps by people with T1D. An advantage of the current study was the use of the wrist-finger flexor muscles. These muscles are less activity dependent than muscles in the leg as the primary physical activity in most people is walking.

A limitation of the present study is that T1D subjects were adults with relatively healthy HbA1c levels indicating they had well controlled blood sugar levels. Most of the subjects in our study were 19-22 years of age. While our study did include some adults who were 25-31 years of age, the sample size for this adult population was too low to provide confidence in a sub-group analysis. Future studies should evaluate more subjects who are older or with poor glycemic control or a longer disease duration to determine health implications of a heavier disease burden. A disadvantage of using the forearm is that it does not reflect training adaptations related to different activity levels. Our experiment did not test if T1D impacts the ability to train mitochondria and impact endurance (43).

## CONCLUSIONS

There were no differences seen between well-controlled adults with T1D and similar controls in muscle endurance, mitochondrial capacity, or muscle reperfusion rate. We had hypothesized that T1D might have reduced muscle endurance and mitochondrial capacity due to suggestions of accelerated mitochondrial aging in this population. Poor glycemic control is also associated with vascular pathology. We also did not see evidence within our generally well controlled T1D subjects of relationships between muscle endurance, muscle mitochondrial capacity, and muscle reperfusion rate and HbA1c levels. It is possible that older or less well controlled subjects with T1D would have evidence of impaired muscle function. More testing would need to be performed to understand the effects of T1D muscle endurance, mitochondrial capacity, and microvascular function in this subjects with T1D across ages and with different levels of glycemic control.

## Acknowledgements

The results of this study are presented clearly, honestly, and without fabrication, falsification, or inappropriate data manipulation. This research was supported by the NIH grant 1R41DK112497 and the UGA Center for Undergraduate Research Opportunities Research Assistantship Program (R.A. Hewgley and B.T. Moore). We would like to express gratitude to Hannah Bossie, Jared Brizendine, Melissa Erickson, and Shuan Goh for assistance with NIRS and muscle endurance methods and Peri Levey for her assistance with our protocol and subjects.

